# Evidence for timing in the midsession reversal task with rats in operant conditioning boxes

**DOI:** 10.64898/2026.03.16.712080

**Authors:** Marcelo Bussotti Reyes, Fillipi da Rosa Ferreira, Gabriel Gobbo Felix da Silva, Marcelo Salvador Caetano, Armando Machado

## Abstract

The midsession reversal (MSR) task is frequently used to study behavioral flexibility and decision strategies in animals. In a typical version of the task, subjects complete 80 trials in which they choose between two simultaneously presented stimuli, S1 and S2. During the first 40 trials, responses to S1 are reinforced, whereas responses to S2 are not. The contingencies then reverse without warning: From trial 41 to 80, only responses to S2 are reinforced. In birds, performance in this task is often characterized by anticipatory and perseverative errors around the reversal point, suggesting a reliance on elapsed time since the session began. In contrast, rats tested in operant conditioning chambers typically show near-optimal performance with few errors, a pattern often interpreted as evidence that rats rely primarily on local reinforcement cues rather than temporal information. The present study investigated whether rats exclusively rely on local cues in the MSR task or whether temporal information also contributes to the decision process. Two groups of rats were trained with different intertrial intervals (ITIs; 5 s or 10 s) while the reversal point remained fixed at Trial 41. During acquisition, both groups diplayed similar learning rates and near-optimal steady-state performance with minimal anticipatory or perseverative errors. However, when the ITI was manipulated in probe sessions, systematic shifts in switching behavior emerged. Rats adjusted their choices according to the temporal midpoint experienced during training rather than the nominal trial number of the reversal. These results suggest that rats rely on a mixed strategy that integrates local reinforcement cues with global timing information. Temporal control may therefore be present even when it is not expressed during standard training conditions.

## Introduction

Behavioral flexibility has been extensively studied with animals to elucidate the neural and cognitive mechanisms underlying adaptive control and decision-making. One experimental paradigm designed to examine this flexibility is the Mid-Session Reversal (MSR) task, originally introduced by Cook and Rosen (2010). Pigeons were trained in a matching-to-sample procedure in which after pecking at a sample stimulus (red or green central key), they chose between two comparison stimuli (red and green side keys). During the first half of the session, the pigeons received food for choosing the comparison that matched the sample; during the second half of the session, they received food for choosing the comparison that differed from the sample. That is, the reinforcement contingencies reversed abruptly at midsession without any signaling cue. How animals adjust their choice in response to, or in anticipation of the contingency reversal was the main issue under investigation.

Subsequently, Rayburn-Reeves et al. (R. Rayburn-Reeves et al., 2011) simplified the task considerably by substituting the conditional discrimination involved in the match–to-sample procedure by a simultaneous discrimination with two stimuli, S1 and S2 (red and green key). In 80-trial sessions, reinforcement followed S1 choices during the first 40 trials and S2 choices during the last 40 trials. A constant interval (ITI) separated consecutive trials. To characterize steady-state performance, both studies examined the probability of choosing S1 as a function of trial number, and both found a characteristic ogive-like psychometric curve with indifference close to the session midpoint (Fig. 1A).

**Figure 1.**
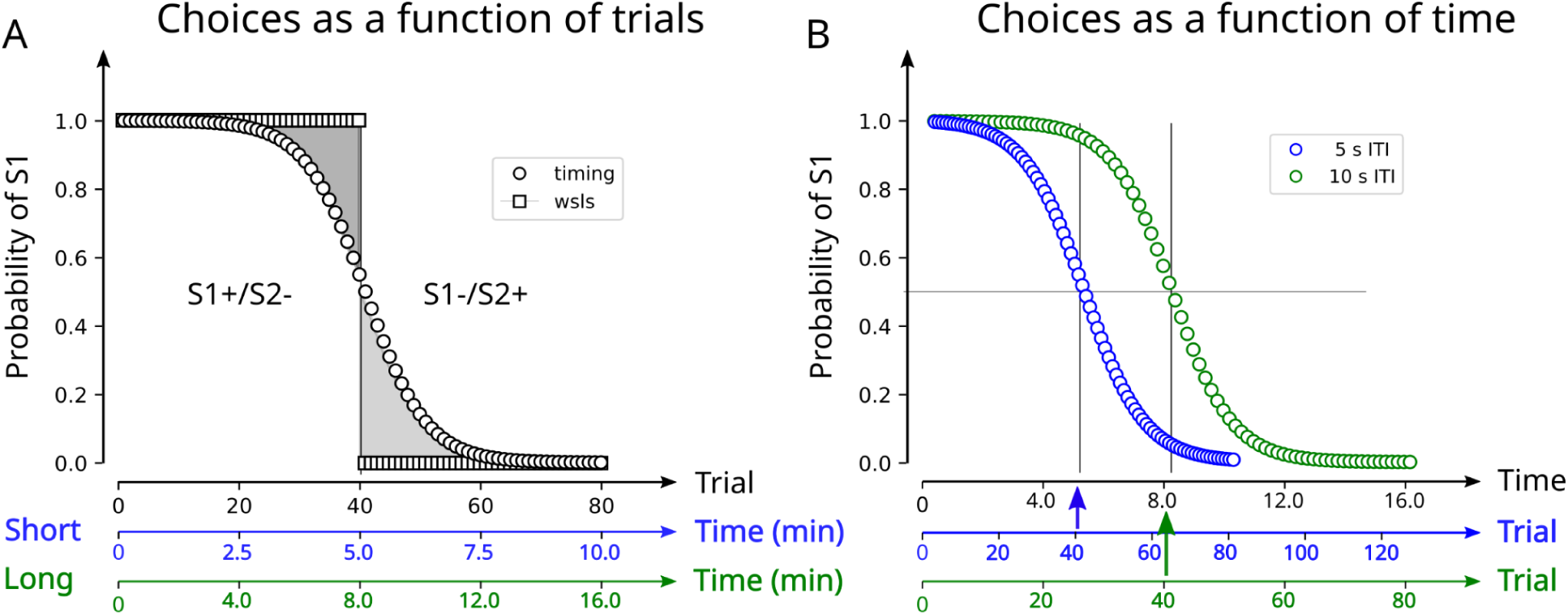
Schematics of idealized behavioral patterns or strategies in the midsession reversal task. (A) Probability of choosing S1 (P_S1_) as a function of trial number for two distinct strategies. Squares represent a win–stay/lose–shift strategy wherein the subject chooses S1 until the first non-reinforced trial and then switches to S2 for the remainder of the session, collecting all reinforcers except one. Circles represent a time-based strategy, in which elapsed time since session onset serves as the cue to switch preferences from S1 to S2 around midsession. This strategy produces anticipatory errors (premature choices of S2 before trial 40; early decline of the curve) and perseverative errors (continued choices of S1 after trial 41; slow post-reversal decay). The colored lines below the graph indicate the temporal progression for two groups trained with different intertrial intervals (ITIs): Short (5 s, blue) and Long (10 s, green). Although the trial-based psychometric functions for the two groups overlap, the same functions plotted across time do not because the trials proceed at different speeds the Short group reaches the reversal trial in ∼5 min, whereas the Long group reaches it in ∼8 min. (B) The time-based strategies shown in A (ogive-like overlapping functions) are replotted against time into the session, showing that the time-based psychometric functions no longer overlap.

Extensive research on the MSR task with pigeons (Carvalho et al., 2019; Cook & Rosen, 2010, Laude et al., 2014; McMillan & Roberts, 2012; R. Rayburn-Reeves et al., 2013; R. M. Rayburn-Reeves et al., 2011; Santos et al., 2019; Soares et al., 2020; Stagner et al., 2013) and with starlings (Machado et al., 2023; Salinas et al., 2025) surprisingly, but consistently, demonstrated that these animals exhibit anticipatory and perseverative errors around the reversal point, revealing a strategy relying on the time elapsed from the beginning of the session (as represented in Fig. 1A, circles). Although sometimes pigeons exhibit performance approaching optimal levels (Laude et al., 2014; McMillan & Roberts, 2012), avian behavior in the MSR task is generally characterized by a substantial number of errors. In contrast, studies with other species, such as monkeys (R. M. Rayburn-Reeves et al., 2017) and humans (R. M. Rayburn-Reeves et al., 2011), report minimal evidence of anticipatory and perseverative errors when tested in operant paradigms. Instead, these animals predominantly use the outcome of the last trial to guide behavior on the next trial: If the choice was rewarded, repeat it; otherwise, change it. Applied consistently, this win-stay/lose-shift strategy outperforms the use of temporal cues because it ensures the collection of all available rewards except the one on the reversal trial.

A few studies investigated rats’ performance in the MSR task, showing that they employ varied strategies. Depending on training conditions, rats either perform near optimally by relying largely on immediate feedback (local cues) or utilize timing/counting strategies (global cues), leading to anticipatory and perseverative errors. When trained in a T-maze, rats make errors comparable to those made by pigeons (McMillan et al., 2014). In contrast, rats trained in operant conditioning chambers produce few errors (R. Rayburn-Reeves et al., 2013), resembling performance patterns observed in monkeys and humans. However, even in operant chambers, rats do produce anticipatory and perseverative errors, although infrequently. The presence of these errors suggests that, contrary to earlier assumptions (R. M. Rayburn-Reeves et al., 2018), rats in operant chambers may rely in part on global cues. In other words, the rats may have learned multiple cues, global and local, but the local cues may dominate or overshadow the global cues during training. Moreover, revealing potential learning of global cues may require appropriate test conditions.

The present study investigated the extent to which rats utilize global (time or counting) cues in the MSR task by systematically manipulating intertrial intervals (ITI, Fig. 1B). We trained two groups of rats: one with a 10 s ITI and the other with a 5 s ITI. Our results showed no differences between groups during acquisition at the steady state; the rats produced very few anticipatory and perseverative errors, consistent with previous studies that reported little evidence for global cues. However, when we then changed the ITIs in probe sessions – while maintaining the reversal at a constant trial number (trial 41) – marked differences emerged between the two groups. Rats trained with shorter ITIs shifted their choices from S1 to S2 significantly earlier, anticipating the reversal point well before trial 41. Conversely, rats trained with longer ITIs delayed shifting preference from S1 to S2 until several trials after trial 41. Notably, both groups reversed preference according to the temporal midpoint experienced during training, the moment of the reversal trial, indicating reliance on timing cues. Using three distinct test structures, we show that rats in the MSR task employ a mixed strategy, integrating timing-based global cues with local outcome-based cues (win-stay/lose-shift).

## Methods

### Animals

Fourteen male Wistar rats participated in the study. They were about 4 months old and weighed around 300 g at the beginning of the experiment. We obtained the animals from the Center of Pharmacology and Molecular Biology (INFAR) from the Universidade Federal de São Paulo. All procedures were previously approved by the Animals Care and Use Committee of the Universidade Federal do ABC (protocol number 2358161122). Upon arrival at the laboratory, the rats were randomly assigned to one of two experimental groups (7 rats per group). They were weighed and handled daily for approximately 5 minutes during one week before starting the experiments. They received food daily after the experiments, in an amount sufficient to maintain them at 85-90% of their ad-libitum weight, measured at the beginning of the handling and weighing procedures.

### Apparatus

The experiment used six chambers from Med-Associates^@^, each 25cm wide, 30cm high, and 30cm deep. Each chamber was equipped with a central magazine pellet dispenser that delivered 45 mg sucrose pellets (Schraibmann LTDA, Brazil) into a food cup, two non-retractable levers located at the right and left sides of the pellet dispenser, two light bulbs placed above each lever which provided diffuse illumination of approximately 200 lx, and a houselight on the top of the back wall. We programmed the behavioral tasks using MedPC^@^ software with a temporal resolution of 2 ms.

### Lever Press Training (Fixed ratio 1)

Rats underwent initial lever-press training over two to four sessions, depending on the animal. The goal was to habituate rats to the chamber and establish consistent lever pressing for food. Sessions consisted of illuminating both cue lights above the levers, each one randomly assigned to steady (ON) or flicker (FL). Pressing either lever resulted in immediate delivery of a sugar pellet. Sessions lasted either 40 minutes or until the animal received 60 pellets.

### Intermediate training

During three sessions, only the S1 cue light was presented on each trial (either steady or flickering at 1 Hz, 500 ms on/off), a press on the corresponding lever delivered a reward, and a press on the other lever had no scheduled consequences. The next three sessions introduced an ITI of 5 or 10 s (according to group), during which all lights were off, and any lever press reset the ITI timer, postponing the onset of the next trial. Training continued until two sessions were completed, with more than 70 correct responses out of 80 trials. Following these initial phases, rats were trained with different procedures that attempted to train them in the MSR task, which included: changing the flickering frequency from1 Hz to 5 Hz; removing the ITI after incorrect responses; and adding correction trials, i.e., incorrect responses produced the ITI and repeated the previous trial until a correct response occurred. Rats showed great difficulty learning to choose the correct stimulus (steady or flicker) when they were presented randomly at different locations (left or right). Hence, we decided to present S1 and S2 always on the same side of the food cup. Fixating the stimulus greatly improved learning, allowing us to move to the final procedure. Then the experiment proper began.

### Baseline

Once a day at 3 pm, the rats underwent a training session comprising 80 trials. Each trial started with the presentation of the two stimuli (ON and FL), henceforth designated as S1 and S2, counterbalanced across rats. The stimuli always appeared on the same side, S1 on the left and S2 on the right, for half of the rats, and with the opposite mapping for the remaining rats. A press in either lever turned both lights off. If the choice was correct, the rat received a sugar pellet, and the ITI began; if the choice was incorrect, the ITI began immediately.

Table 1 shows the contingencies during the baseline (Train) sessions for each group: Short, trained with a 5 s ITI, and Long, trained with a 10 s ITI. Responses during the ITI reset the timer (r in the Table). The standard MSR reinforcement contingencies applied for both groups: choices of S1 but not of S2 were reinforced from trials 1 to 40, and choices of S2 but not S1 were reinforced from trials 41 to 80. Training lasted 25 sessions.

**Table 1:** Contingencies for each group during Training and Testing. Group Short trained with a 5 s resetting (r) ITI for 80 trials. The standard MSR reinforcement contingencies applied. In Test 1, the ITI was 10 s and resetting; the reinforcement contingencies remained as in Train. Test 2 was like Test 1, except the 10 s ITI did not reset (nr). Test 3 was like Test 2, except that S2 was reinforced from trials 21 to 80. Group Long trained with a 10 s resetting ITI for 80 trials, and the standard MSR reinforcement contingencies. In Test 1, the number of trials increased to 160, the 5 s ITI was resetting, and the contingencies rewarded S1 on all trials and S2 from trials 41 to 160. Test 2 was like Test 1, except for a non-resetting ITI. Test 3 was like Test 2, except that S1 was reinforced only from trials 1 to 80.

| Experimental Contingencies |  |  |  |  |
| --- | --- | --- | --- | --- |
| Group Short |  |  |  |  |
|  | Train | Test 1 | Test 2 | Test 3 |
| Trials | 80 | 80 | 80 | 80 |
| ITI(s) | 5, r | 10, r | 10, nr | 10, nr |
| S1+ | 1-40 | 1-40 | 1-40 | 1-40 |
| S2+ | 41-80 | 41-80 | 41-80 | 21-80 |

Table 1: Contingencies for each group during Training and Testing.
|  | Group Long |  |  |  |
| --- | --- | --- | --- | --- |
|  | Train | Test 1 | Test 2 | Test 3 |
| Trial | 80 | 160 | 160 | 160 |
| ITI(s) | 10, r | 5, r | 5, nr | 5, nr |
| S1+ | 1-40 | 1-160 | 1-160 | 1-80 |
| S2+ | 41-80 | 41-160 | 41-160 | 41-160 |

#### Test 1

For both groups, the ITI changed during test sessions, to 10 s for Group Short and to 5 s for Group Long. All other procedural details remained as in Baseline sessions, except for the following for Group Long: the total number of trials increased to 160, with S1 reinforced throughout all 160 trials, and S2 reinforced from trials 41 to 160. The rationale for this change concerns the behavior predicted from Group Long during the test session. If they adopt a pure time-based strategy, they should persevere with S1 well after trial 40, perhaps until trial 80. The range of trials in which any choice could be rewarded aimed to inhibit the intrusion of local cues before rats could exhibit a time-based behavior (see Machado et al., 2023, Soares et al., 2024).

#### Test 2

This test replicated Test 1 except that responding during the ITI did not reset the timer (nr in Table 1).

#### Test 3

Replicated Test 2, but with different reinforcement contingencies. In Group Short, S1 was reinforced from trials 1 to 40, and S2 was reinforced from trials 21 to 80. For Group Long, S1 was reinforced from trials 1 to 80, and S2 was reinforced from trials 41 to 160.

After each test, the animals received four retraining sessions in which all procedural details, including the ITI,were identical to baseline. To illustrate, a rat from Group Short experienced in succession three conditions, each with a 5-s ITI during training and a 10-s ITI during testing: Condition 1, Train and Test 1; Condition 2, Train and Test 2; and Condition 3, Train and Test 3.

### Retraining with different ITI

After completing the three train/test conditions, the animals switched groups, i.e., rats from Group Short (5 s ITI) trained with long ITI (10 s) and vice versa, for a total of 26 sessions. The training with an ITI and the testing with the other ITI progressed as in the first phase, for a total of three Conditions (Train and Test 1, Train and Test 2 and Train and Test 3). We combined the data from the two phases.

### Data description

To assess the effect of shifting from training to testing, we compared data from each test session with data from the three preceding training sessions, which served as a baseline. We averaged the animals’ performance across the two phases, i.e., we disregarded the phase in our analysis.

**Table 2:** Sessions used for data analysis in each Condition (Training and Test) and in each Phase.

| Condition | Phase 1 |  |  |  | Phase 2 |  |  |  |
| --- | --- | --- | --- | --- | --- | --- | --- | --- |
|  | Training |  |  | Test | Training |  |  | Test |
| 1 | 22 | 23 | 24 | 25 | 43 | 44 | 45 | 46 |
| 2 | 27 | 28 | 29 | 30 | 48 | 49 | 50 | 51 |
| 3 | 32 | 33 | 34 | 35 | 53 | 54 | 55 | 56 |

### Data Analysis

Behavioral responses were analyzed in terms of the probability of responding to S1 (P_S1_) as a function of either trial number or session time. For trial-based analyses, we first averaged the responses across the three baseline sessions of each Condition, and then across animals to obtain the average training function and the average test function. Error bands represent the 95% bootstrap percentile-based confidence intervals – for each session trial, 1000 samples with replacement were collected from the individual psychometric function.

For time-based analyses, we aggregated responses using 1-minute bins (e.g., 0–1 min, 1–2 min). For each bin, we averaged P_S1_ across the three baseline sessions for each animal, averaged across rats the training sessions and the test sessions to obtain the group average time-based psychometric functions. The 95% percentile-based bootstrap confidence intervals were computed for each time bin in the same way they were computed for each trial. ITo assess changes during test sessions, we compared the group-average psychometric function from each test with the average of the three preceding training sessions. Specifically, we first computed the mean P_S1_ per rat and per condition (training or test), either across trials or time bins, yielding one psychometric function per rat per condition. We then averaged across rats. Error bars and shaded areas in figures reflect the 95% confidence interval of the group mean, computed for each trial or time bin across the 14 average baseline sessions and the 14 single-test sessions.

To estimate the psychophysical parameters of interest (point of subjective equality [PSE] and difference limen [DL]), we performed logistic regression analyses at the trial level for each animal and session, including test and baseline sessions. The binary response variable was the outcome of each trial, coded as 1 (S1) or 0 (S2), and the predictor variable was the trial number. The baseline parameters were computed for each rat and session, then averaged across baseline sessions for each rat to compare with the test sessions. For the figures, we average across all rats for each condition.

To summarize group performance, PSE and DL values were averaged across animals for each condition and group (train and test, short and long). We used these averages in final plots and group-level comparisons. Logistic regressions were performed independently for analyses using trial number and session time as the predictors.

## Results

Rats required approximately 20 sessions to achieve steady performance (Fig. 2A), with no differences between groups in learning rate or asymptotic accuracy (circa 92%). After rats reached steady performance, we investigated how the probability of choosing S1 evolved during the 9 training sessions used as a baseline. Rats in both groups made a few errors (Fig. 2B): The psychometric functions overlapped, slightly decreasing as the reversal trial approached, and then somewhat abruptly, in a step-like manner, after the reversal trial. The results suggest that rats relied mostly on a win-stay/lose-shift strategy. When plotted as a function of time (Fig 2C), the same data reveal the different time evolution between groups, with Group Short typically experiencing the reversal trial approximately 5 min into the session. In contrast, Group Long experienced the reversal trial approximately 9 min into the session. It is noteworthy that the rats progress differently in the trial rate (trials per minute). Such variation stems from differences in choice latencies (the delay between the stimulus presentation and the response) and the number of responses during the ITI (that causes a timer reset, see methods). These individual differences shape the average psychometric functions into a smooth sigmoid-like curve, while the individual rats tend to switch to S2 more abruptly. Also, these smooth average curves are scale transforms of each other, as the overlap of the trial-based curves (Fig. 2B) implies.

**Figure 2:**
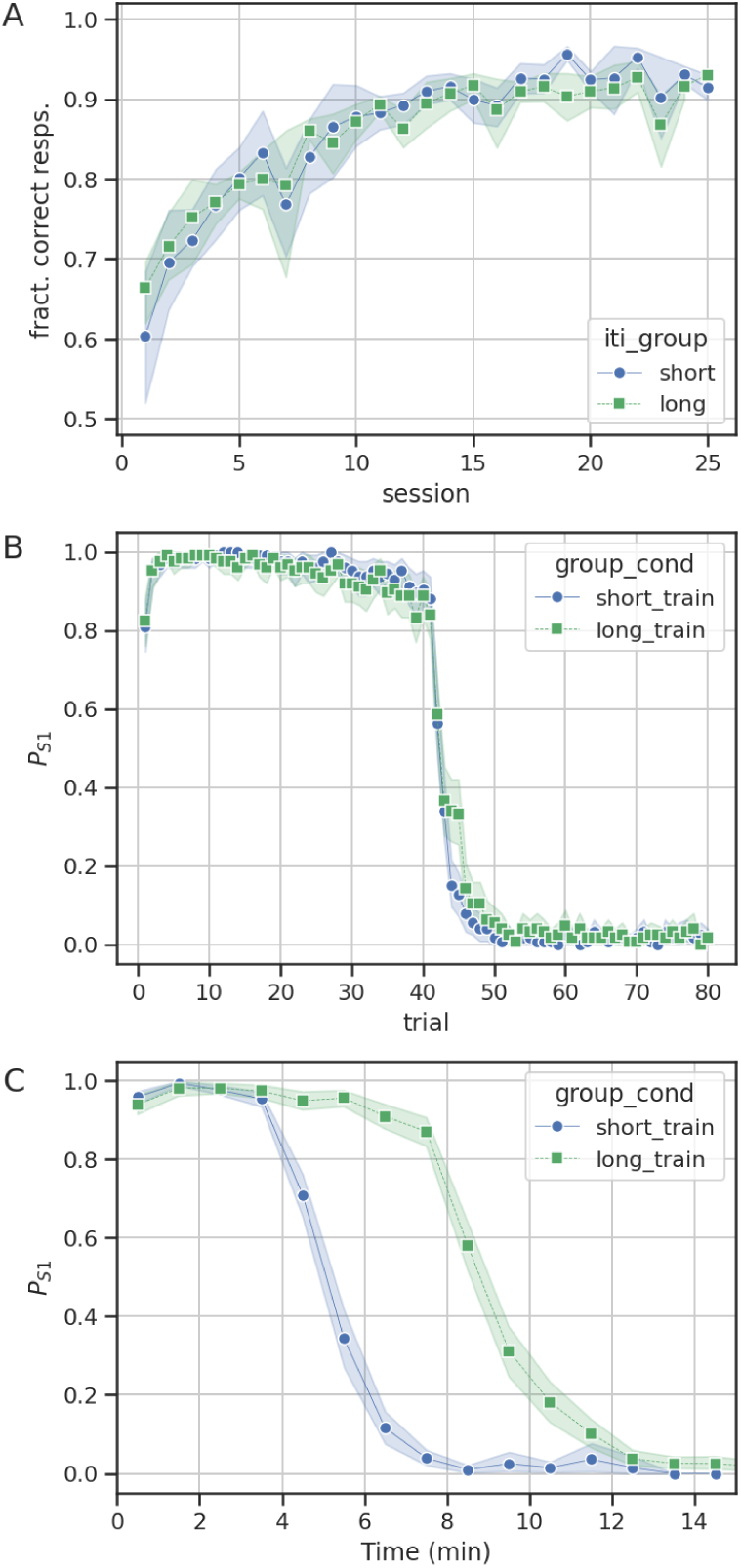
A) Fraction of trials reinforced during the acquisition phase separated by group (Short or Long) showing that the groups evolved similarly. B) and C) Performances of both groups (short and long) during training, averaged over all the 9 baseline sessions. B) Probability of responding S1 as a function of the trial. Rats typically respond to S1 almost entirely before the reversion (trial 41), then start choosing S2, as shown by an abrupt decline in the curve. By trial 50, virtually all rats in both groups had chosen only S2. C) Same data as in B, but now showing the probability of choosing S1 (averaged over 1 min bins) as a function of time, showing that the time of transition from S1 to S2 differs between groups due to the different inter-trial intervals.

### Test 1: Switching the ITI duration between groups

Even though in baseline sessions rats showed little anticipation, when we manipulated the ITI, responses from both groups markedly deviated from baseline and in opposite directions, suggesting that animals relied – at least partially – on a timing strategy (Fig 3A, see supplementary Figs. 1 and 2 for individual data). Group Short, with 5 s ITI in Training (blue curve), experienced 10 s ITI in testing, which means that the session pace approximately halved. In this case, as the orange curve shows, the rats began choosing S2 much earlier than during baseline. Notice that only S1 choices led to reinforcement before trial 40. Hence, the animals were clearly weighing their choices using a cue other than the trial outcomes. However, the average progression of P_S1_ did not follow the same shape as the baseline, showing a plateau from trials 20 to 40, followed by a decay after trial 40. The plateau suggests that other processes influenced the rats’ choices, possibly the repeated exposure to non-reinforced choices of S2 (extinction) before the reversal.

**Figure 3:**
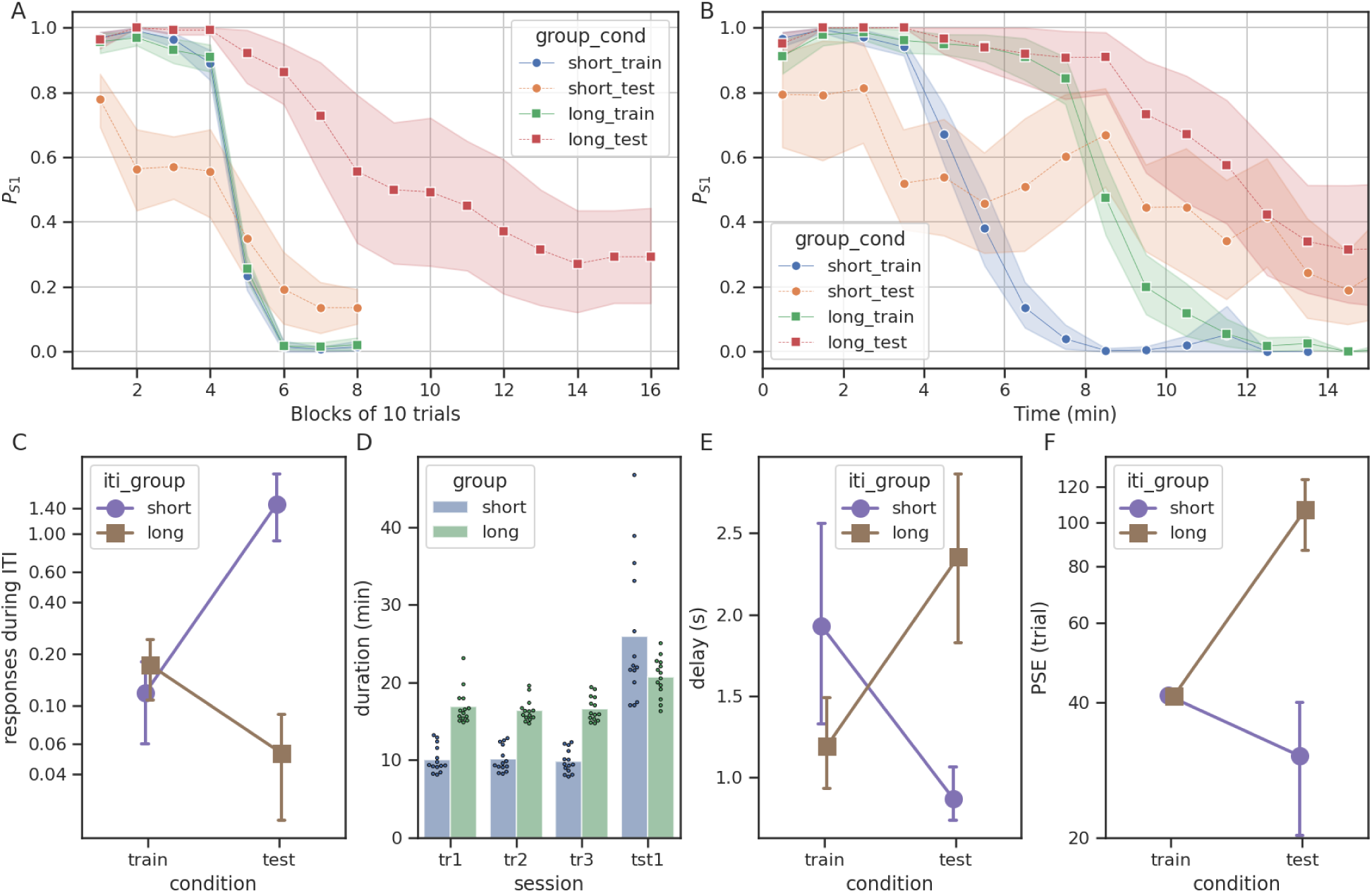
Results from Test 1. A) Probability of responding S1 as a function of trials (in blocks of 10). The blue (Group Short, 5 s ITI) and green (Group Long, 10 s ITI) curves show the performance in the three sessions preceding the test. The shaded areas represent 95% bootstrap confidence intervals for the mean across all 14 rats. The orange and red curves show Group Short and Long test responses, respectively. B) Same results as in A, but with the probability of S1 responses plotted as a function of time for the first 15 minutes of the session. Responses were aggregated into 1-min time bins (see methods). Shaded areas show the 95% confidence intervals for the mean of 14 rats per group. C) Average number of responses (in log scale) during the ITI for each group in train and test. D) Duration of the three training and test sessions. The test sessions for both groups were expected to last about the same as the training sessions of Group Long, but 5 rats for Group Short had very long sessions due to increased ITI responding (which reset the ITI timer). E) Mean latency to press the lever for each group in train and test. Error bars represent 95% CI of the mean across rats. F) Mean trial-based point of subjective equality (PSE, in log scale) obtained in the four Conditions (train and test for groups Short and Long). Error bars represent 95% CI of the mean across rats.

Group Long, with 10 s ITI in training (green curve), experienced 5 s ITI in testing. hence, a session progression twice as fast. The probability of choosing S1 decreased more slowly across trials (red curve) than in the baseline. Because S1 was reinforced across all 160 trials, there was no reason to switch to S2 unless rats used cues other than choice outcomes. Again, animals clearly did not adopt a pure WSLS strategy: all rats began to choose S2 before the end of the session.

We also investigated how P_S1_ evolved over time (rather than across trials; Fig. 3B). If rats relied solely on time to choose the lever, the curves for each group in the train and test conditions should superimpose. Clearly, they did not, supporting that rats relied on cues distinct from timing to make their choices.

When testing Group Short (orange curve), the plateau observed in Fig. 3A (cf. orange curve between blocks 2 and 4) turned out to be a valley in time (Fig. 3B), with rats sampling S2 earlier than in baseline (≈3.5 min into the session), and then choosing the two levers almost equally on average. If time were the only factor, there would be no reason for such an increase in S2 choices.

The overall disruption in preference was probably also due to a salient difference in the effective ITI durations between train and test sessions. In all sessions, responses during the ITI were punished by resetting the ITI timer. Rats in Group Short were trained with a 5 s ITI and therefore learned to refrain from responding during that period (Fig. 3C). However, this learning did not generalize to the longer 10 s ITI used during Test 1. These rats produced significantly more premature responses (Fig. 3C), which increased the overall session duration (Fig. 3D). In contrast, Group Long was not affected by the ITI manipulation: Accustomed to longer 10 s ITIs during training, they did not produce significantly different numbers of premature responses when they were tested with the 5 s ITI.

The foregoing ITI-related behavior may help to explain how response latencies, i.e., the time between the onset of the stimulus lights and the lever press, changed in test sessions compared to training sessions (Fig. 3E). The mean latency in Group Short decreased perhaps because these rats were expecting a much shorter ITI and hence were probably ready to respond when the lights turned on. The mean latency in Group Long increased for the opposite reason: because they did not expect the shorter ITI, rats were more likely to be far from the levers when the lights turned on.

Finally, we investigated how individual rats transitioned to S2 using the point of subjective equality (PSE) obtained from the logistic regression (Fig. 3F). The results show a different response from each group when tested, strengthening the idea that rats from Group Short started choosing S2 earlier (smaller trial-based PSE) and rats from Group Long later (larger trial-based PSE) than in baseline. A two-way repeated measures ANOVA revealed significant main effects of Group (*F*(1, 13) = 40.39, *p* = 2.52×10⁻⁵, η²□ = 0.42) and Condition (Train vs. Test, *F*(1, 13) = 27.56, *p* = 1.57×10⁻⁴, η²□ = 0.22), as well as a significant interaction between Group and Condition (*F*(1, 13) = 39.48, *p* = 2.82×10⁻⁵, η²□ = 0.43).

### Test 2: Keeping the reversal time more reliable

The results from Test 1 suggest that rats also rely on timing when choosing the lever in the MSR task. Still, the variability in reversal time during the test session in Group Short added a confounding variable, making it difficult to interpret the strategy rats in this group used in the test. To address this issue, Test 2 (Fig. 4, see supplementary Fig. 3 and 4 for individual data) eliminated the reset contingency following ITI responses. This adjustment allowed rats to experience more uniform ITI durations and reduced the variability in the reversal time across subjects (Fig. 4D).

**Figure 4:**
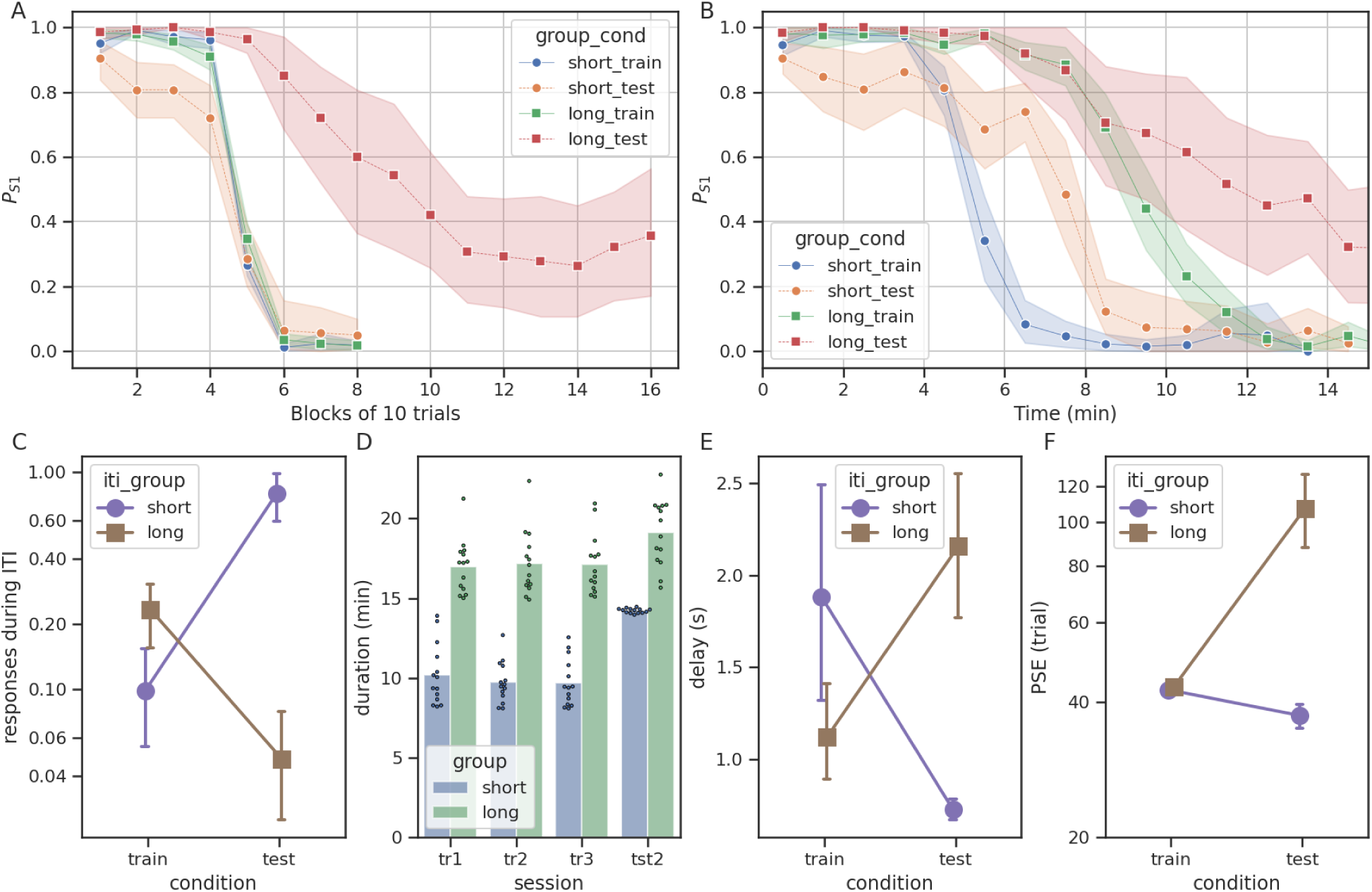
Results from Test 2. A) Probability of choosing S1 as a function of the trials for Test 2. In this case, the only difference from Test 1 was that the timer was not reset when animals responded during the ITI, thereby decreasing variability in reversal times. For the Short group, the tendency to choose S1 earlier in trials remained comparable to Test 1, resulting in a (narrower) plateau between blocks 2 and 3. However, P_S1_ decreases to near zero faster after the transition than in the same curve in Fig. 3. In Group Long, the slower decay of P_S1_ remains as in Test 1 (Fig. 3). B) Same data as in A, but showing P_S1_ as a function of time in bins of 1 min. C) Average responses (in log scale) during the ITI for each group in baseline and test. D) Individual data of the session duration for the three training (tr) sessions before Test 2 (tst2). E) Average delay to press the lever for each group in baseline and test. F) Average PSE (in log scale) for all conditions.

The results largely replicated the patterns observed in Test 1: rats in Group Short (tested with longer ITIs) again exhibited an earlier decline in P_S1_, whereas rats in Group Long (tested with shorter ITIs) reversed their preference much later than trial 41 (Fig. 4A). Crucially, however, the P_S1_ curve for Group Short showed a steeper decrease following trial 40, similar to baseline, with narrow confidence intervals at and after the reversal trial. Thus, with a consistent ITI duration, the rats were more likely to encounter the reversal point under similar temporal conditions, thereby reducing between-subject variability.

When we plot the same against elapsed time (Fig. 4B), they confirm that timing, while influential, was not the sole factor guiding the rats’ choices. Consider the difference between the two groups in the first half of the test session. In Group Long (red curve in Fig. 4B), the rats seem to follow a temporal cue from the beginning to approximately 8 min into the session, with the testing and baseline functions overlapping considerably. Afterward, the two functions diverge, as preference for S1 does not decrease as fast as during baseline. In Group Short, the testing (orange) and baseline (green) curves never overlap, with preference for S1 declining considerably later in testing. Time cues appear to exert a stronger influence on the long-to-short than on the short-to-long ITI changes.

Toward the end of the test sessions (Fig. 4B, final segments of orange and red curves), the two groups exhibited markedly different asymptotic behavior. In Group Short, all rats reliably transitioned to choosing S2 after the reversal and maintained this preference for the remainder of the session, yielding P_S1_ values close to zero. In contrast, rats in Group Long showed more variable behavior. The probability of responding to S1 never declined completely to zero, even well after the reversal point. This persistence indicates that some rats continued to choose S1 intermittently throughout the session.

A closer inspection of individual behavior supports this interpretation: among the fourteen rats in Group Long, one chose S1 on all 160 trials, five switched completely to S2, and the remaining eight alternated between S1 and S2 (see supplementary material). In Group Short, 10 of the 14 rats produced only S2 responses on the last 10 test trials.

This between-group asymmetry is likely driven by differences in the reinforcement contingencies during testing. In Group Long, both S1 and S2 responses were reinforced from trials 41 to 160. As a result, although the timing cue might have prompted a transition around trial 80, continuous reinforcement of S1 likely interfered with a full preference shift. Thus, the P_S1_ curve failing to reach zero in Group Long may reflect the influence of local outcome-based cues overriding global time-based cues.

Interestingly, a complementary effect may have occurred in Group Short. Based on time cues, the rats in this group were expected to reverse their preference near trial 20, but S2 choices were not reinforced before trial 41. The outcome cues may have counteracted the time cues until the reversal trial. In this case, the absence of reward following early S2 choices delayed the reversal of preference guided by timing cues. In contrast, in Group Long, the same delay was observed due to the presence of a reward following late S1 choices. To summarize, our data suggest that in both groups, local outcome-based and global time-based cues competed to influence choice.

Regarding premature responses (Fig. 4c), the rats followed a similar pattern from Test 1, with a slight decrease in Group Long and a strong increase in Group Short. For Group Short, however, the increase in responses was not as large as in Test 1, most likely because the ITI timer did not reset in Test 2. Hence, even if the rats responded prematurely, the ITI remained approximately the same - double the ITI in the training sessions - and, since it did not reset, the opportunities to produce premature responses decreased compared to Test 1. The regularity in the evolution of the test session is also evident in the session duration (Fig. 4d), where all the rats in Group Short finished the session in about 14 min, with little variation between rats (column tst2, blue). Except for this, the duration of the sessions for the baseline for both groups and the test for Group Long remained similar to Test 1 (Fig. 3d). Complementary to these results, the delay (latency) to respond (Fig. 4e) when the stimuli turned on also present opposite effects, with the reduction of the latency for Group Long and increase in Group Short. The pattern was virtually identical to that in Test 1.

The analysis of the PSE (Fig. 4F) revealed that, whereas the mean PSE increased significantly from training to testing in Group Long, it decreased slightly in Group Short. The two-way ANOVA revealed significant main effects of group (F(1,13) = 47.24, p = 1.1 × 10⁻⁵, η²□ = 0.78), condition (F(1,13) = 30.58, p = 9.7 × 10⁻⁵, η²□ = 0.70), and the interaction between ITI and condition (F(1,13) = 45.46, p = 1.4 × 10⁻⁵, η²□ = 0.78)

### Test 3: Aligning Task Structure and Reinforcement Contingencies Across Groups

Results from Tests 1 and 2 indicated that rats were strongly influenced by the task pace, relying on elapsed time as a cue for switching behavior. They also suggested that local reinforcement contingencies play an important role in guiding choices. However, an asymmetry persisted between groups: rats in Group Short delayed switching until S2 responses were reinforced, whereas rats in Group Long often failed to switch completely, most likely because S1 continued to be reinforced. We designed a test to remove the asymmetry between groups and further examine the role of local reinforcement in modulating behavior. We intended to make the reinforcement contingencies similar, allowing the two groups to follow temporal cues without counteracting their effects with local cues that either penalized early transitions to S2 or reinforced S1 for too many trials. In Test 3, for Group Short, S1 was reinforced from trials 1 to 40 (as in training), while S2 was reinforced from trial 21 to 80 – ensuring that temporally appropriate early switches to S2 would be rewarded. For Group Long, S1 was reinforced from trials 1 to 80 – avoiding reinforcement of S1 beyond the expected reversal point – and S2 from trials 41 to 160.

The results once again confirmed a clear difference between groups, driven by a strong influence of temporal cues (Fig. 5A, see supplementary Fig. 5 and 6 for individual data). Additionally, both groups exhibited behavioral patterns distinct from those observed in Test 2. On average, rats in Group Short (orange) showed an earlier transition to S2, with PS1 decreasing to approximately 65% in block 3 and a marked decline from blocks 2 to 3. In Group Long (red), rats now transitioned fully to S2, as expected, with virtually no S1 responses after block 10. This complete transition confirms that the persistence of S1 responses in Test 2 was due to continued reinforcement of S1 beyond the reversal point.

**Figure 5:**
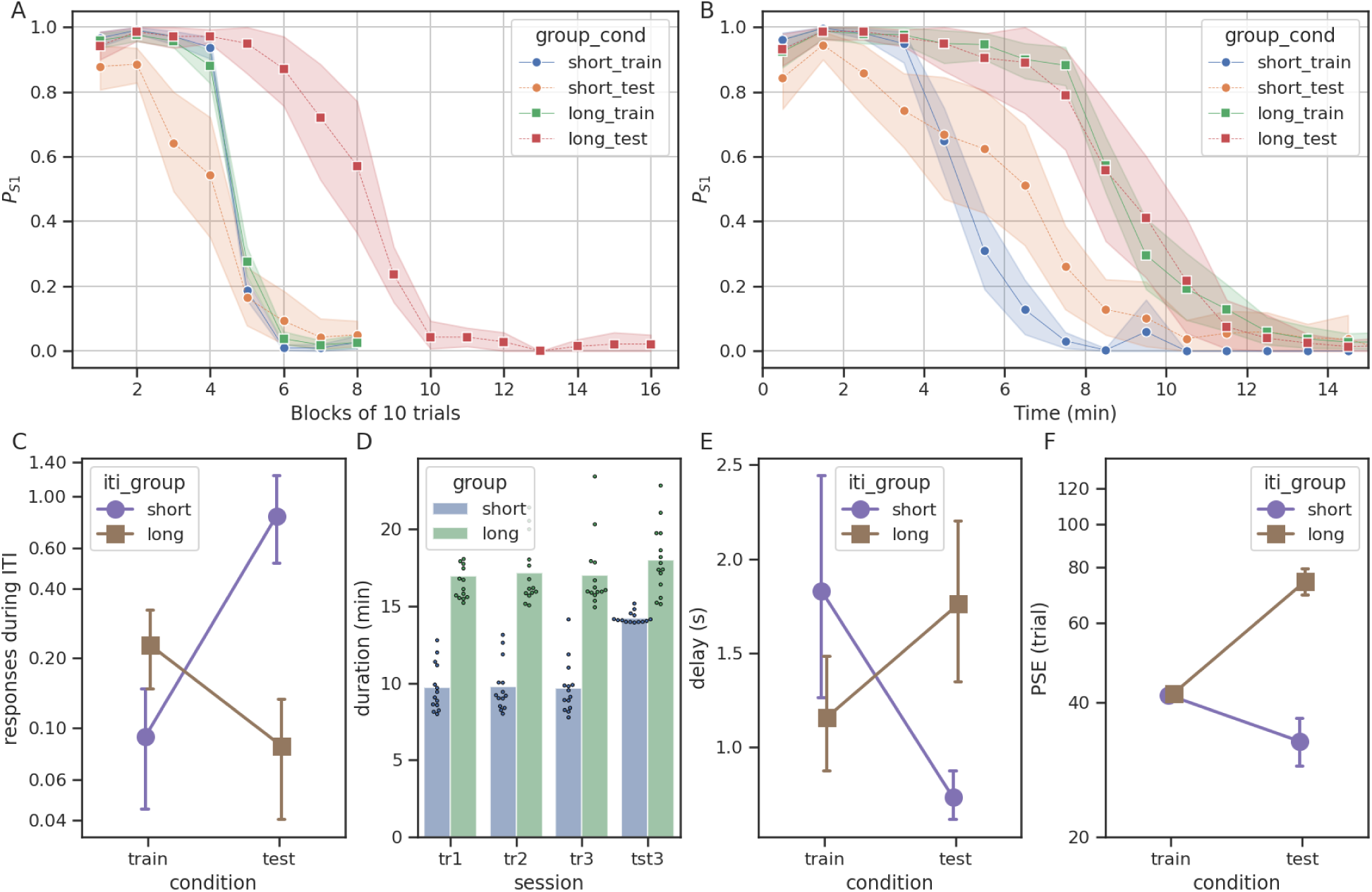
Results for Test 3. A) Probability of S1 as a function of the trials (average over blocks of 10). When tested, Group Short (orange line) had three phases of reinforcement: S1+ from trials 1 to 20, S1+S2+ from trials 21 to 40, and S2+ from trials 41 to 80. The Group Long also had three phases: S1+ from trials 1 to 40, S1+S2+ from trials 41 to 80, and S2+ from trials 80 to 160. B) Same data as A, but showing P_S1_ as a function of time in bins of 1 min. C) Average responses (in log scale) during the ITI for each group in baseline and test. D) Individual data of the session duration for the three training (tr) sessions before Test 3 (tst3). E) Average delay to press the lever for each group in baseline and test. F) Average PSE (in log scale) for all conditions.

**Figure 6:**
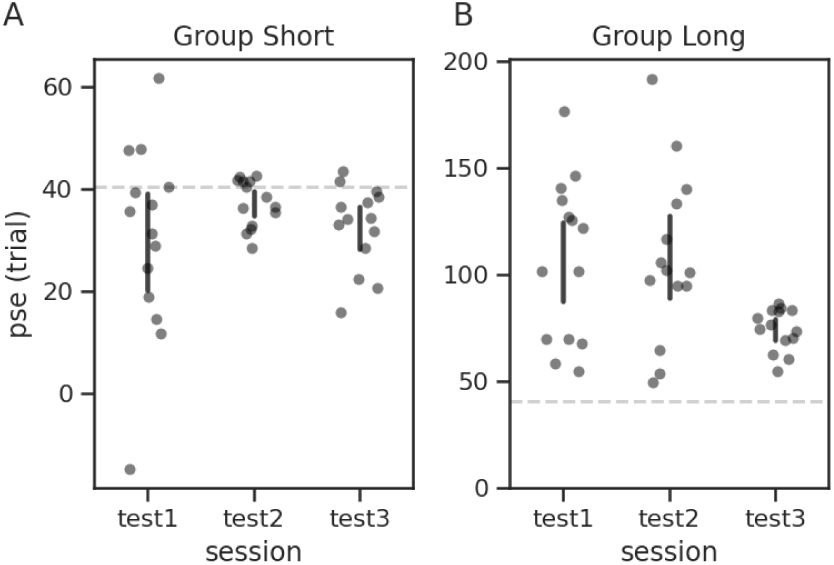
Individual PSE exclusively for the three test sessions. The vertical bars show the 95% confidence interval for the mean, and the dashed horizontal lines show the average PSE across training sessions.

Regarding the number of responses during the ITI, session duration, the average delay to press the levers, and the PSE (Figs. 4c-f), the results remained unchanged from Test 2. There were significant main effects of ITI group (F(1,13) = 293.23, p = 2.7 × 10⁻¹⁰, η²□ = 0.96) and condition (F(1,13) = 35.33, p = 4.9 × 10⁻⁵, η²□ = 0.73), as well as a significant interaction between group and condition (F(1,13) = 251.26, p = 7.0 × 10⁻¹⁰, η²□ = 0.95).

Finally, we compared the point of subjective equality (PSE) across the three test types (Fig. 6). Average PSEs did not differ within the short ITI group (Fig. 6A): repeated-measures ANOVA yielded F(2, 26) = 1.70, p = .20, partial η² = .12. In contrast, a significant effect of test type was observed in the long ITI group (Fig. 6B), F(2, 26) = 7.50, p = .003, partial η² = .37. Bonferroni-corrected pairwise comparisons revealed that PSE values in Test 3 differed significantly from both Test 1, t(13) = 3.40, p = .014 and Test 2, t(13) = 3.43, p = .013, whereas no evidence of difference between Test 1 and Test 2 (t(13) = −0.06, p = 1.00). The difference in Test 3 for Group Long effect was due to the fact that every rat switched before or around trial 80, the point at which S1 ceased to be rewarded. Such a local cue is much more marked than the temporal cues, and hence we expect less variability between individuals (as seen in Fig. 6B, Test 3). Overall, these results confirm the tendency of the rats to persist in S1 for longer than the temporal cues indicate, as long as this choice is rewarded. In Group Short, even when S2 was reinforced after trial 20 (Figure 6A, Test 3), only three of the rats switched around this trial (as expected by the timing hypothesis). In Group Long, a small group of rats switched before trial 70, but many took more than 100 trials to switch, persevering in S1 (Figure 6B, Tests 1 and 2). Only in Test 3, when S1 ceases to reinforce after trial 80, all rats transitioned almost immediately after this event.

## Discussion

In this study, we investigated how rats behave in the midsession reversal task when the choices S1 and S2 are always presented at the same side, i.e., when the discrimination between S1 and S2 is spatial. We trained two groups in this task, differing only in the intertrial interval (ITI). We showed that both groups acquired the task at a comparable speed, with no detectable difference between groups. These results highlight a distinction between rats and pigeons: previous reports support that pigeons perform worse when trained with 5 s ITI, compared to 1.5 s ITI, in a spatial version of the MSR (Laude et al., 2014) analogous to what we use in the present study. Even though we used longer ITIs (5 and 10 s), rats performed similarly to pigeons with 1.5 s --around 95% correct choices for the entire session, on average, after training. If the reduction of the performance with longer ITI results from the limited working memory capacity of preserving the choice made in the last trial, our results show that rats can keep such memories for longer intervals. These results from pigeons also suggest that rats’ performances may decline if they were trained with longer ITI durations.

After training, as in previous reports (R. Rayburn-Reeves et al., 2013; R. M. Rayburn-Reeves et al., 2018; Smith et al., 2016), rats displayed a sharp transition at the reversal point and made few anticipatory or perseverative errors. These results have previously been interpreted as evidence that rats rely predominantly on local cues (e.g., following a win-stay/lose-shift strategy, WSLS). However, our subsequent manipulations of ITI challenged this interpretation. Surprisingly, we found that rats are highly sensitive to changes in the temporal structure of the task: When we modified the ITI – and thus the session tempo –, rats systematically shifted their choices in accordance with elapsed session time, at times prioritizing temporal cues over reinforcement-based cues. The reliance on temporal information emerged even though timing was not explicitly required during training, revealing that rats extract and employ temporal regularities to adjust their choices.

Our study involved three complementary tests. Together, these tests provide evidence that rats use both temporal and local cues to guide their choices and quantify how they weigh these cues. In the first test, we doubled the ITI for Group Short and halved it for Group Long. Given that Group Long would experience twice as many trials (and could face an enormous number of extinction trials), we used slightly different reinforcement contingencies for this group, reinforcing S1 for the entire session (see Machado et al., 2023; Soares et al., 2023 for a discussion of the contingencies during testing). For Group Short, we changed only the ITI, leaving every other aspect of training unchanged. The ITI manipulation produced opposing deviations from baseline. Rats from Group Short pressed S2 more during testing than in baseline before trial 40, suggesting that elapsed time – rather than a pure reward-following strategy like WSLS – also guided the early part of the session. Yet their average PS1 curve plateaued between trials 20–40, indicating that many rats switched back to S1, perhaps because the early S2 presses were unrewarded. Conversely, rats from Group Long, now experiencing a faster session, chose S1 longer than during baseline; only late in the session did they drift toward S2. Estimates of the Point-of-Subjective-Equality (PSE) confirmed the divergence: PSEs were smaller than in baseline for Group Short and larger than in baseline for Group Long, revealing that rats weighed timing cues when deciding which lever to press, but those cues were modulated and sometimes overridden by the reinforcement contingencies.

A key insight from our data is that near-perfect performance (absence of anticipatory and perseverative errors) does not imply the absence of timing or of learning temporal cues. The abrupt transition from S1 to S2 is effectively incompatible with the scalar property of interval timing (Gibbon, 1977), for if rats during baseline sessions relied exclusively on temporal cues, Weber’s law should engender far more anticipatory and perseverative errors than we and others have observed, a number closer to that observed in experiments with pigeons (Cook & Rosen, 2010, p. 201), starlings (Machado et al., 2023) and rats trained in a T-maze (McMillan et al., 2014). From this discrepancy, we can conclude that rats do not deploy a timing strategy in baseline sessions, and that temporal control appears only when temporal and reinforcement cues conflict, as when the ITI changes from training to testing.

Smith et al. (2016) reversed the reinforcement contingencies at a random trial every session, so that the reversal point was unpredictable. They showed that rats produced more anticipation errors when the reversal occurred later in the session. Santos and Sanabria (2020) only partially replicated these results, finding a similar difference only when the probability of reinforcing a correct response was reduced to 50%. In different ways, both studies showed that time can influence rats’ decisions in the MSR task, particularly when the weight of local cues is reduced. Our results add to these findings by showing that, even without random factors (reversal trial, probability of reinforcement), rats still learn about the task’s underlying temporal structure, but they reveal it only when the local and temporal cues “disagree”. Such evidence supports that acquisition itself may be tied to time, since learning involves temporal relations between events (Balsam et al., 2010; Gallistel & Gibbon, 2000).

Our data also shows an asymmetry between the two groups. Rats from Group Short evaded transitioning early in the sessions, even when the local and temporal cues agreed (Test 3). This effect was at odds with that observed in studies with pigeons (McMillan & Roberts, 2012; Soares et al., 2020; Zentall, 2020) and starlings (Machado et al., 2023). In general, birds have produced stronger leftward shifts when tested with a doubled ITI, compatible with the timing hypothesis. On the other hand, the same birds produced more modest rightward shifts with the halved ITIs. Machado et al. (2023) investigated the interference of generalization decrement, arguing that subjects would face a large number of extinction trials when the ITI was halved, suggesting that the test structure might be unfit for the task. Those results led us to continue reinforcing S1 throughout the session (Tests 1 and 2) or until trial 80 (Test 3). For that reason, our data cannot resolve the issue of whether rats in Group Long would produce modest rightward shifts if we had kept the same reinforcement contingencies during testing as during training. However, given the strong effect of stopping rewarding S1 after trial 80 (Test 3, Group Long), and the overall susceptibility of rats to local cues, we hypothesize that rats would produce even shorter rightward shifts than pigeons and starlings.

Together, our results support the view that behavior in the midsession reversal task reflects a dynamic weighting between local cues that subsume the win-stay/lose-shift strategy and global cues that subsume a timing strategy; a mixture rather than a pure strategy. Rats maintain an accurate yet latent sense of elapsed time. Still, they rely on it only when its predictions deviate sufficiently from the regular training conditions and do not conflict with the local cues – conditions created here by altering ITI duration and the S1 reward schedule. By showing that this latent timing signal can be unmasked without introducing randomness, we also support the view that timing might be part of learning and memory in the task. The gap between avian and rodent findings is thereby reduced.

## Acknowledgements

This work was supported by grants from FAPESP (Grant 2022/16315-0) and CNPq (INCT NeuroComp 408389/2024-9) to MBR. AM was supported by the Portuguese Foundation for Science and Technology (FCT 2022.04861.PTDC and UIDB/04810/2020). MSC and AM were supported by the Brazilian National Institute of Science and Technology on Behavior, Cognition, and Teaching (FAPESP Grant 2026/01540-9; and CNPq Grant 409051/2024-1) and by FAPESP (Grant 2022/00312-1). We also thank the Timing and Cognition Lab at UFABC for useful discussions and feedback. The authors used artificial intelligence-based language tools exclusively to improve the clarity and readability of parts of the text that had already been written by the authors.

## Supplementary Material

### Individual data from Experiment 1, Test 1

#### Individual data for Test 1

**Supplementary Figure 1:**
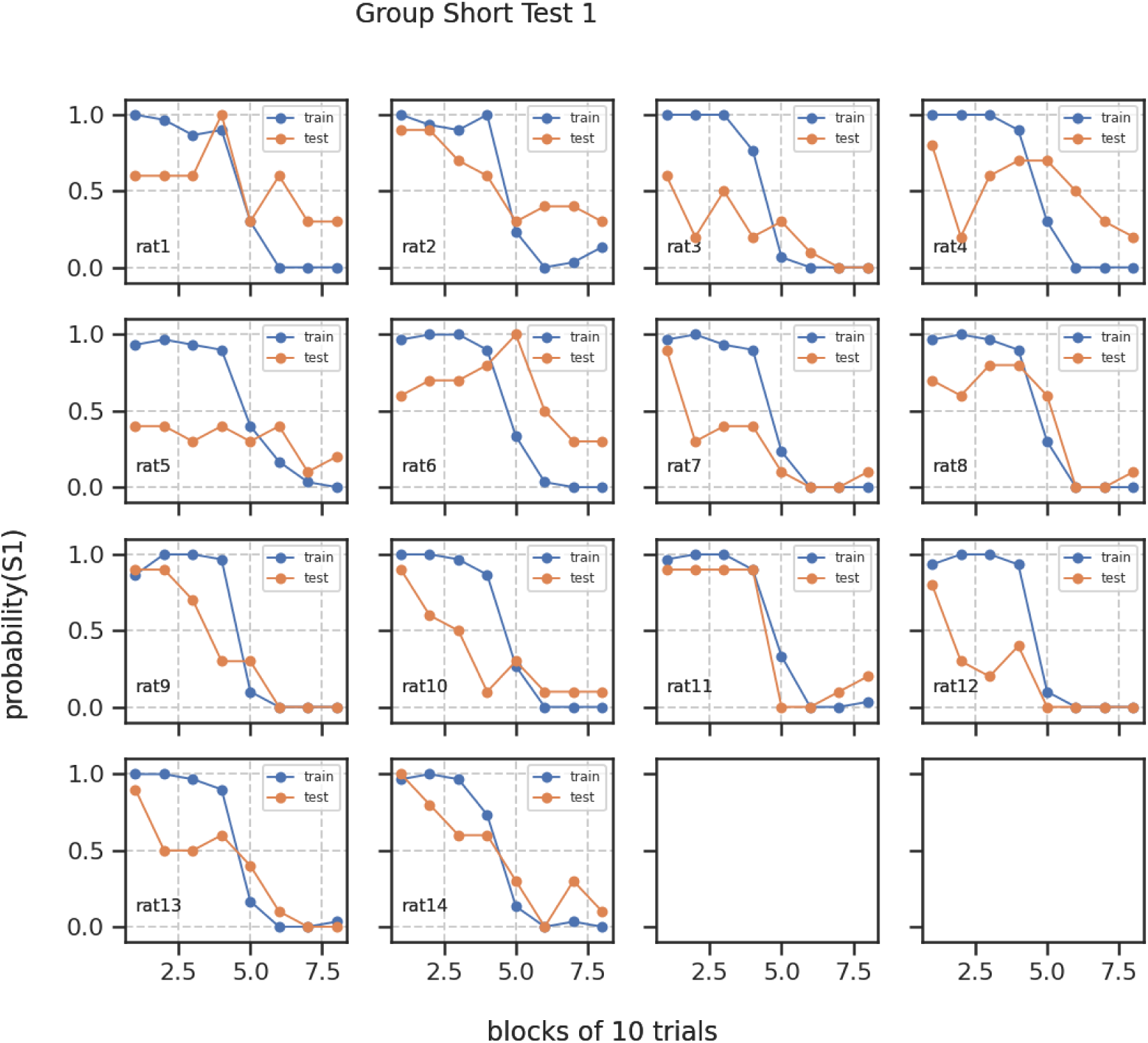
Individual responses in Test 1 for Group Short.

**Supplementary Figure 2:**
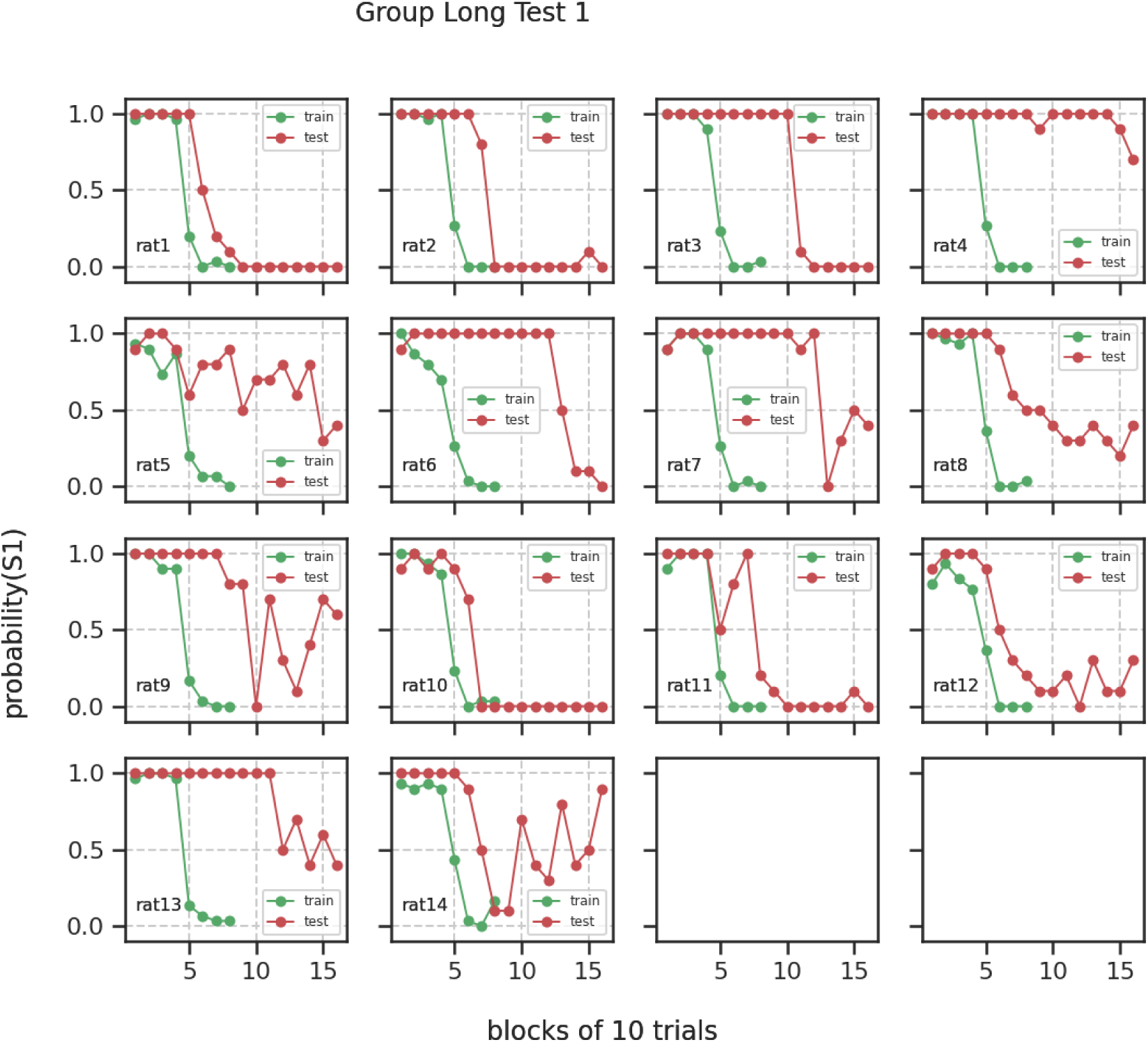
Individual responses in Test 1 for Group Long.

#### Individual data for Test 2

**Supplementary Figure 3:**
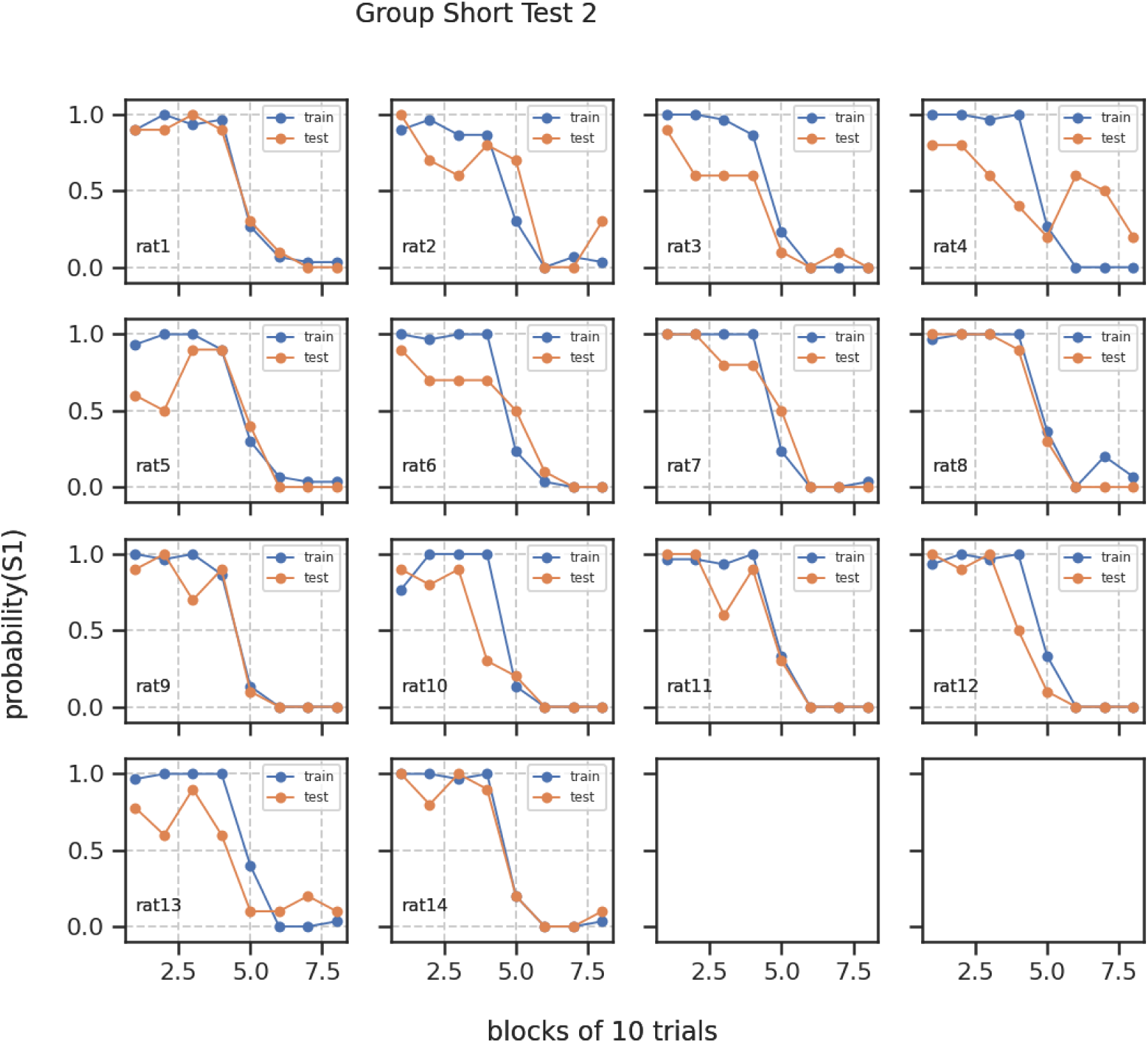
Individual responses in Test 2 for Group Short.

**Supplementary Figure 4:**
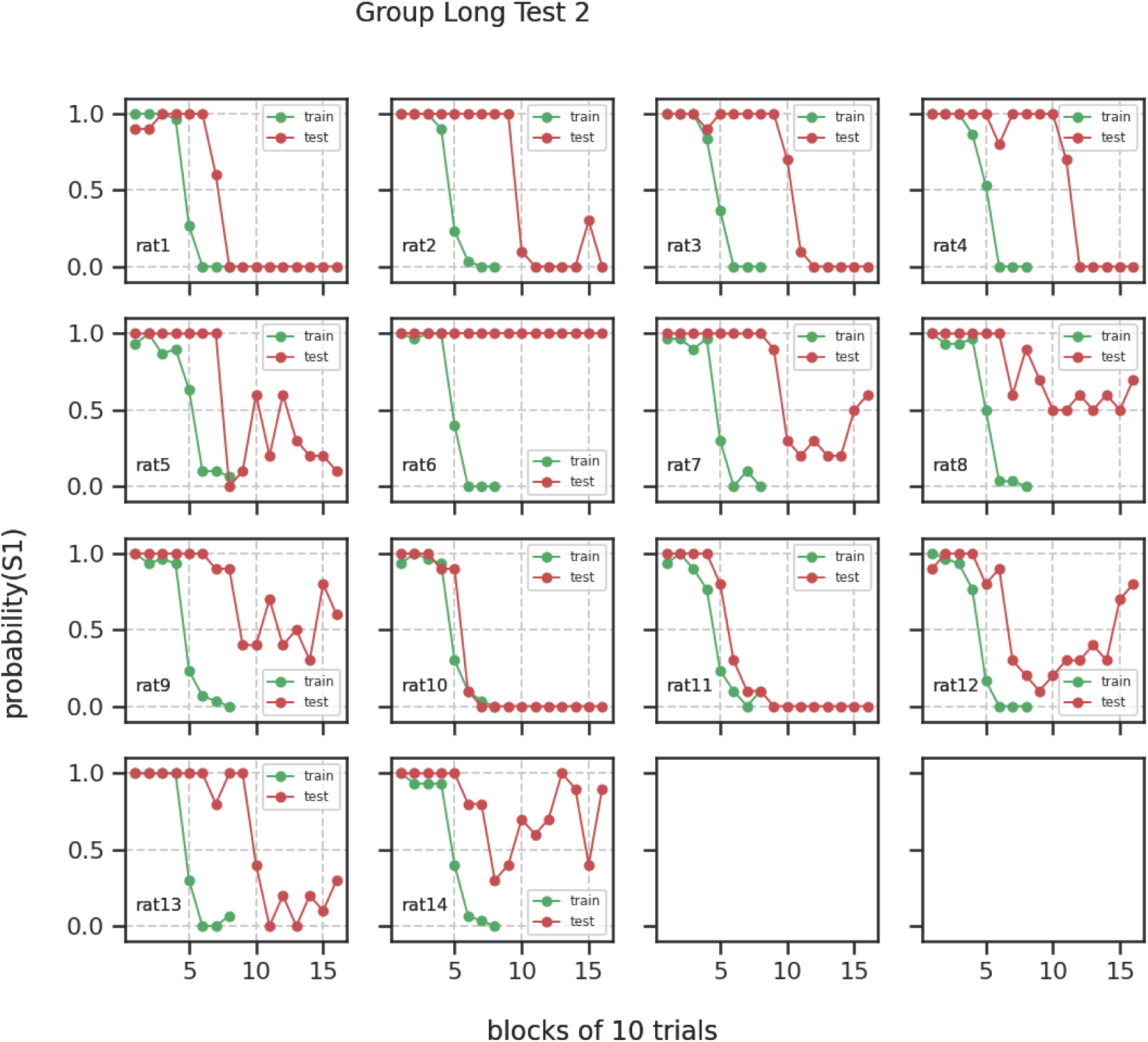
Individual responses in Test 2 for Group Long

#### Individual data for Test 3

**Supplementary Figure 5:**
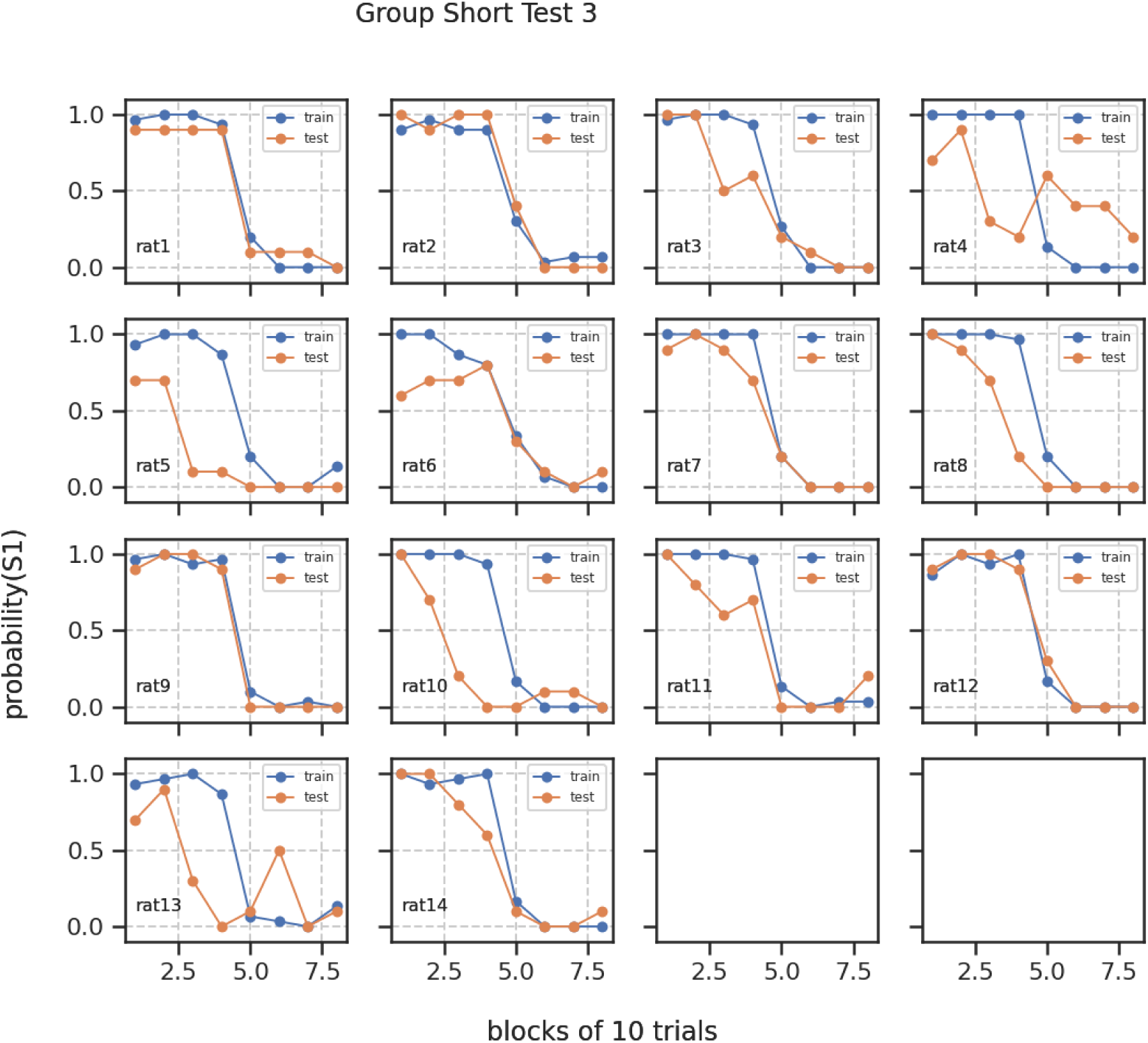
Individual responses in Test 3 for Group Short

**Supplementary Figure 6:**
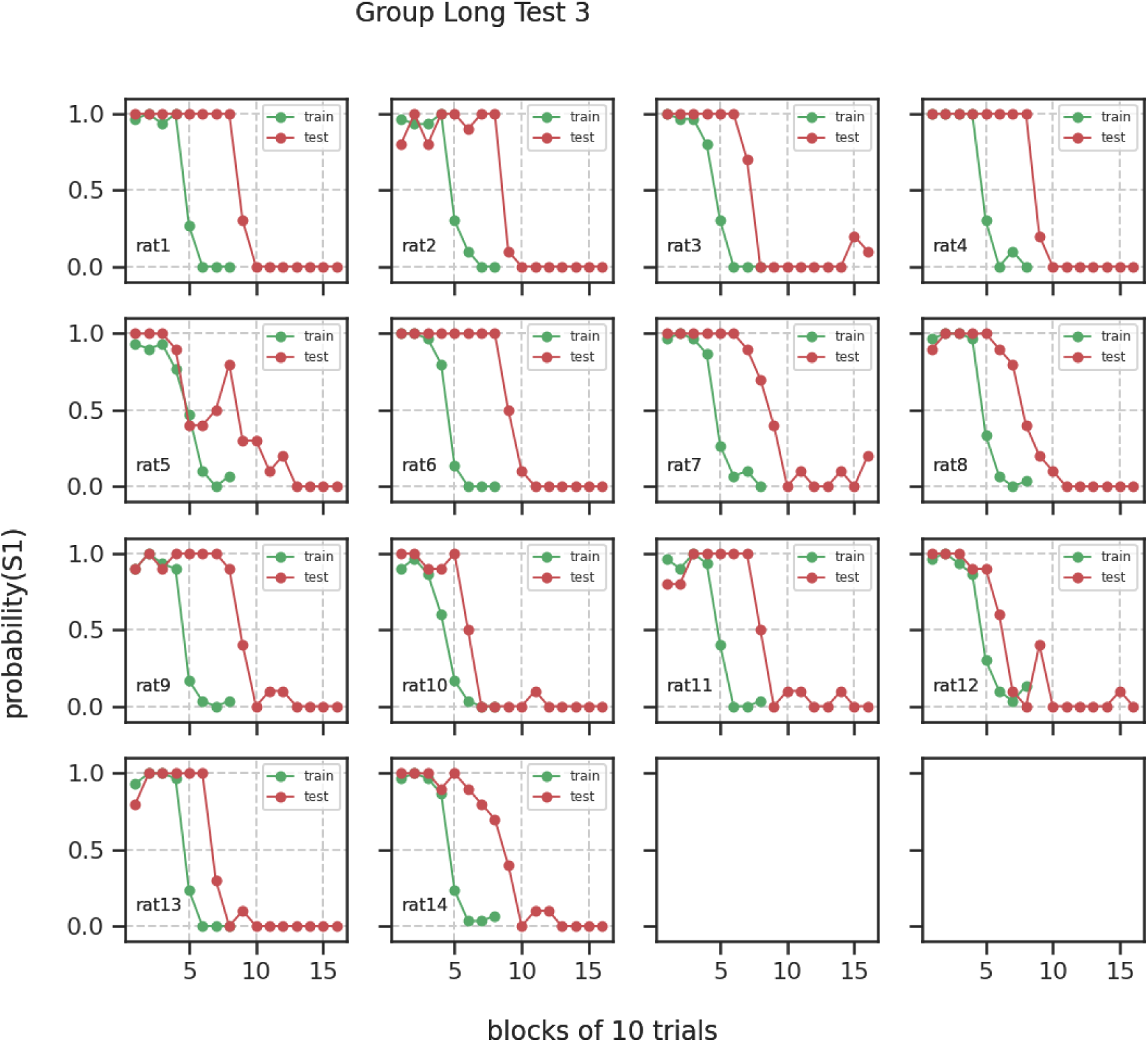
Individual responses in Test 3 for Group Long

